# Tissue-expression profiles unveils the gene interaction of hepatopancreas, eyestalk, and ovary in precocious female Chinese mitten crab, *Eriocheir sinensis*

**DOI:** 10.1101/420182

**Authors:** Xiaowen Chen, Jun Wang, Xin Hou, Wucheng Yue, Shu Huang, Chenghui Wang

**Affiliations:** Key Laboratory of Freshwater Aquatic Genetic Resources, Ministry of Agriculture, Shanghai, 201306, China; National Demonstration Center for Experimental Fisheries Science Education (Shanghai Ocean University), Shanghai, 201306, China; Shanghai Engineering Research Center of Aquaculture, Shanghai, 201306, China

**Author notes:** Corresponding author: Chenghui Wang Address: 999, Hucheng huan Road, Lingang New City, Shanghai, 201306, China. These authors contributed equally to this study (Xiaowen Chen, Jun Wang).

**Keywords:** transcriptome, ovary development, genetic network

## Abstract

Sexual precocity is a serious and common biological phenomenon in animal species. Large amount of precocity individuals was identified in Chinese mitten crab, *Eriocheir sinensis*, which caused huge economical loss every year. However, the underlying genetic basis of precocity in *E. sinensis* is still lack. In this study, histology observation, comparative transcriptome was conducted among different stages of precocious one-year old and normal two-year old *E. sinensis*, tissue-expression profiles of ovary, hepatopancreas, and eyestalk tissues were presented and compared. Genes associated with lipid metabolic process, lipid transport, vitelline membrane formation, vitelline synthesis and neuropeptide hormone related genes were upregulated in the ovary, hepatopancreas and eyestalk of precocious *E. sinensis*. Our results indicated eyestalk involved in neuroendocrine system providing neuropeptide hormone that may induce vitellogenesis in hepatopancreas and further stimulate ovary development. Hepatopancreas is a site for energy storage, vitellogenin synthesis and may assist to induce oogenesis through lipid transport in precocious *E. sinensis*. The genetic basis of precocity in *E. sinensis* is an integrated gene regulatory network of eyestalk, hepatopancreas, and ovary tissues. Our study provides effective convenient phenotype measurement method for identification of potential precocious *E. sinensis* detection, and valuable genetic resources and novel insights into the research of molecular mechanism of precocity in *E. sinensis*.

## Introduction

Sexual precocity refers to the reproductive system (gonad) that abnormally develop into sexual mature stage in advance during puberty, which is a natural phenomenon in most animal species, even in human [1-3]. Growth and development retardation, increased illness rate, and other associated physiological defects are believed to be the consequences of sexual precocity [1, 4]. Sexual precocity is a complex physiological process that induced by extrinsic environment factors and intrinsic genetic factors [2, 5]. The early development of gonads is considered to be the molecular response to environment factors, such as hormone, nutrition, temperature, disease and so on [5]. The scientific communities are struggled with the genetic mechanism of sexual precocity; however, the underlying mechanism is still elusive.

In vertebrate, the gonad development is believed to be regulated by hypothalamus-pituitary-gonad axis (HPG), however, in invertebrate, the regulation of reproductive system is vague [6, 7]. Chinese mitten crab *Eriocheir sinensis*, an economical crustacean widely cultured in China is also suffered from severe precocious problems [8]. Huge economic losses in the *E. sinensis* aquaculture industry is caused by substantial proportion of precocious *E. sinensis* individuals every year [2, 9]. Environment factors such as temperature, salinity, light, and stocking density are believed to induce sexual precocity in *E. sinensis*, however, the intrinsic molecular feedback to the stimulation of environment factors is largely unknown in *E. sinensis* [10-12].

A similar X-organ-sinus gland complex neuroendocrine system existed in eyestalk is believed to play analogous roles as HPG axis in crustaceans [7, 13, 14]. Gonad inhibiting hormone (*GIH*), molt-inhibiting hormone (*MIH*), crustacean hyperglycaemic hormone (*CHH*), neuropeptide F (*NPF*) genes/neuropeptides expressed and synthesized in eyestalk are believed to play essential roles in regulating gonad development of *E. sinensis* [15]. Meanwhile, hepatopancreas is an essential organ for energy metabolism, providing essential energy source for gonad development of *E. sinensis* [16]. Studies have also indicated exogenous vitellogenin is synthesized in hepatopancreas and transferred to ovary during vitellogenesis process in *E.sinensis* [17]. However, how the environment factors stimulate and activate the gonad early development and what is the specific biological function of eyestalk and hepatopancreas in regulating gonad development was largely unknown in precocious *E. sinensis*.

*E. sinensis* is a catadromous species and the life cycle is generally two years. The mating and spawning occurs during winter in brackish water and the fertilized eggs develop into larvae in spring, then the larva will migrate to freshwater rivers/lakes and spend nearly two years with nearly 20-times molting before they get sexual mature [8, 18]. Generally, the gonad development of *E. sinensis* initiates at the second year. As for precocious *E. sinensis*, the gonad starts to develop and get completely sexual mature in the first year [8]. Molting and growth is terminated in sexual mature precocious *E. sinensis* and these individuals are usually discarded due to their unworthiness in the aquaculture [9]. The most direct and accurate way to identify precocious *E. sinensis* is through histology observation which is quite inconvenient during aquaculture for farmers. During the aquaculture process, precocious female *E. sinensis* individuals are always identified based on the shape of abdominal sternite by experienced farmers (**Figure 1A**) [15, 19]. The female *E. sinensis* gets sexual mature when the abdominal sternite completely covers the whole abdomen (Figure 1A). Previous studies have indicated the ratio of abdominal sternite length is a candidate phenotype character to discriminate the precocious level in female *E. sinensis* [19] (Figure 1A). However, researches link the abdominal sternite length to ovary development stages are limited.

**Figure 1.**
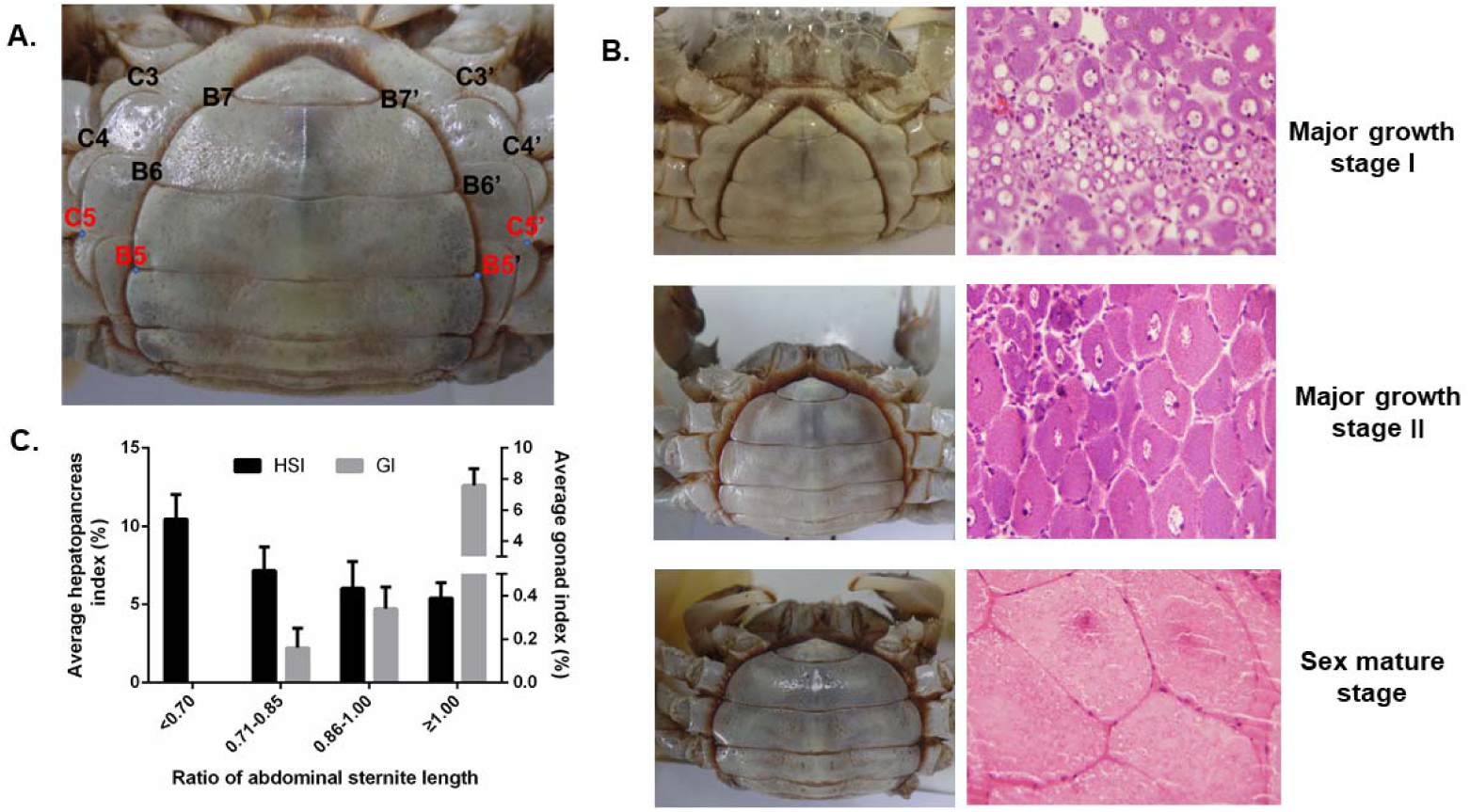
Phenotype characters, histology observation, and hepatopancreas and gonad indexes among different ovary developmental stages of precocious *E.sinensis*. A. Shape of abdominal sternite of female *E.sinensis*. B5-B5’ indicated the outside length of the fifth abdominal sternite, C5-C5’ indicated the inside length of the fifth abdominal sternite. B. Different shapes of abdominal sternite corresponding to different ovary developmental stages in female *E.sinensis*. C. Average hepatopancreas and gonad index among different groups of precocious *E.sinensis.* HIS: hepapancreas index, GI: gonad index.

In this study, potential precocious female *E. sinensis* were firstly collected based on phenotype characters (ratio of abdominal sternite length) and then confirmed by histology observation. Comparative transcriptome was conducted on eyestalk, hepatopancreas, and ovary tissues of precocious female *E. sinensis* from different ovary development stages to 1) provide a convenient way to identify precocious *E.sinensis* in aquaculture, 2) reveal the tissue-expression profiles in precocious female *E. sinensis* in chronological order, 3) identify potential candidate genes/pathways involved in early ovary development, and 4) provide novel insights into the genetic network of eyestalk and hepatopancreas in regulating ovary development in *E. sinensis*.

## Materials and methods

### Animal sampling and ethics

This study was approved by the Institutional Animal Care and Use Committee (IACUS) of Shanghai Ocean University (Shanghai, China). Sampling procedures complied with the guideline of IACUS on the care and use of animals for scientific purposes. All the *E. sinensis* individuals in this study were collected from the Aquatic Animal Germplasm Station of Shanghai Ocean University (Shanghai, China). In this study, potential precocious female *E. sinensis* individuals were firstly collected according to the ratio of abdominal sternite length (B5-B5’/C5-C5’) and then confirmed by histology observation (**Figure 1**).

### Measurement of phenotype characters and histology observation

For the collected *E. sinensis* individuals, phenotype characters such as body weight, ovary weight and hepatopancreas weight were measured, and abdominal sternite length were also recorded according to previous research [19]. Hepatopancreas index (HSI), gonad index (GI), and ratio of abdominal sternite length were calculated according to the following formula.

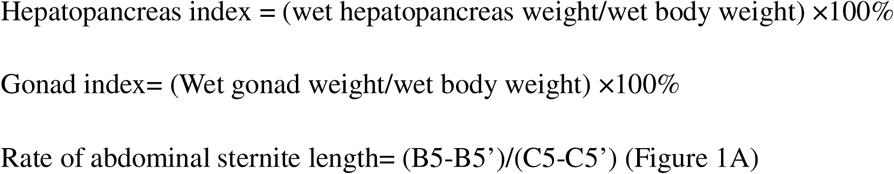

Ovary tissues from potential precocious *E. sinensis* individuals were fixed using Bouin’s fixative (Sangon Biotech, China) at room temperature for 24h. Then the ovary tissue-slices were prepared and stained according to hematoxylin-eosin (HE) staining method. The tissue-slices were observed under a DM500 microscope system (LEIKA, Germany) and Image Analysis Software Toup View.

### RNA isolation and transcriptome sequencing

According to the histology observation results, precocious *E. sinensis* individuals with ovary developmental stages in major growth stage I, major growth stage II, and sexual mature stage were collected with three biological replicates in each group. Meanwhile, three normal developed two-year old female *E. sinensis* individuals (sexual mature stage) were also sampled as control group. All the collected *E. sinensis* were anaesthetized on ice and eyestalk, hepatopancreas, and ovary tissues were quickly collected and immediately stored in liquid nitrogen before RNA extraction. Total RNA was extracted from each collected *E. sinensis* individual with RNAiso Reagent (Takara, China) according to the manufacturer’s instructions. The RNA integrity and quantity were examined using agarose gel electrophoresis and an Agilent 2100 Bioanalyzer (Agilent, Shanghai, China), respectively. A total of 5 μg RNA with an RNA integrity number (RIN) exceeding 8.0 was used for RNA-seq library construction using the Truseq^™^ RNA sample Prep Kit for Illumina (Illumina, USA). These indexed libraries were sequenced on an Illumina Hiseq^™^4000, with 150 bp pair-end reads produced.

### Differential expression and GO enrichment analysis

After sequencing, raw sequencing reads were firstly trimmed using Trimmomatic software[20]. And then, clean reads were mapped to our previously assembled reference transcriptome assembly (NCBI TSA accession number: GGQO00000000) using Bowtie 1.0.0 [21]. Gene abundance, the TPM (transcripts per million transcripts) value was measured using the RSEM 1.3.0 software [22]. The resulting data matrix with expression value (TPM) for all the samples was generated and used as input data for differential expression analysis. Then the differentially expressed genes (DEGs) was measured by DESeq2 software using *P* < 0.001 for the false discovery rate (FDR) and a fold change > 2^2^ [23]. After normalizing the DEG TPM values using log2 and mean centered, cluster analysis was performed using the hierarchical cluster method based on the euclidean distance using heatmap module in R. GO enrichment analysis of the DEGs was conducted using TopGO software with an adjusted *P* value < 0.001 [24]. Pearson correlation was calculated and plotted by corrplot package in R.

### qRT-PCR validation

Quantitative real-time PCR (qRT-PCR) was carried out to validate the DEGs identified in this study. Eight DEGs in ovary, hepatopancreas, and eyestalk were chosen for qRT-PCR assays. PCR primers were designed according to our previous reference transcriptome assembly (**Table S1**). In this study, three reference genes *ubiquitin conjugating enzyme (Ube), beta-actin (*β*-actin),* and *ribosomal S27 fusion protein (S27)* were selected to normalize the gene expression level. qRT-PCR was conducted using SYBR Green Premix Ex Taq (Takara, China) in a QIAxcel real-time PCR system (Qiagen, German). A standard curve was firstly generated to assess amplification accuracy, and primers with an amplification efficiency between 95% and 105%, and Pearson correlation (R^2^) >0.98 were chosen for following qRT-PCR experiments. Three biological and three technical replicates were chosen for each selected DEG. The relative expression was estimated using the 2^−ΔΔCt^ method with normal developed sex mature *E. sinensis* individuals as a calibration control [25]. Relative expression results were presented as the fold-change relative to normal developed sex mature *E. sinensis* individuals. Statistical significance (*P* < 0.05) was determined using one-way ANOVA tests under SPSS 25.0.

### Data archive

Sequencing reads are available at NCBI SRA database (SRR7777398, SRR7777399, SRR7777400, SRR7777401, SRR7777402, SRR7777403, SRR7777404, SRR7777405, SRR7777406, SRR7777407, SRR7777408, SRR7777409)

## Results

### Phenotype characters measurement and ovary histology observation

According to the ratio of abdominal sternite length and the histological observation from collected potential precocious *E. sinensis* individuals, four groups of *E. sinensis* with different ovary developmental stage were clearly identified. Group I, the ratio was less than 0.7 and no clearly ovary tissue was discovered; Group II, the ratio was ranged from 0.71-0.85, the ovary developmental stage was in major growth stage I; Group III, the ratio was 0.86-1.0 and the ovary developmental stage was in major growth stage II; Group IV, the ratio was greater than 1.0 and the ovary stage was completely mature with clearly oocytes (Table 1, Figure 1B). The hepatopancreas index decreased from 10.45% to 5.40% and the gonad index increased from 0 to 7.59% accompanied with the abdominal sternite length ratio increased from 0.70 to 1.00. (**Table 1, Figure 1**).

**Table 1.** Information on phenotype characters and histology observation of precocious *E.sinensis*.

| Rate of abdominal sternite length | No. of individuals | Average weight | Average hepatopancreas index | Average gonad index | Ovary developmental stage |
| --- | --- | --- | --- | --- | --- |
| <0.70 | 6 | 5.62±0.7 | 10.45±1.56% | 0 | Not applicable |
|  |  | 2g |  |  |  |
| 0.71-0.85 | 9 | 13.72±1. | 7.16±1.52% | 0.16±0.0 | Major growth |
|  |  | 23g |  | 9% | stage I |
| 0.86-0.99 | 9 | 19.47±4. | 6.03±1.70% | 0.34±0.1 | Major growth |
|  |  | 01g |  | 0% | stage II |
| ≥1.0 | 6 | 40.90±1. | 5.40±0.98% | 7.59±1.0 | Mature stage |
|  |  | 68g |  | 7% |  |

### Tissue gene expression profiles in precocious *E. sinensis*

Regarding ovary tissue, a total of 957 DEGs were identified among different groups of precocious *E. sinensis*. Three clusters were defined based on the hierarchical clustering results revealing different expression patterns in the ovary of precocious *E. sinensis*. GO enrichment analysis indicated that genes in cluster 1 including sodium-dependent nutrient amino acid transporter 1 (*NAAT1*), MFS-type transporter (*SLC18B1*), solute carrier family 13 member 3 (*SLC13A3*), solute carrier family 10 member 6 (*SLC10A6*), monocarboxylate transporter 12 (*SLC16A12*), glucose-6-phosphate exchanger (*SLC37A4*), low-density lipoprotein receptor 1 (*LDLR-A*), nose resistant to fluoxetine protein 6 (*NRF-6*) genes associated with transmembrane transport for amino acid, creatine, glucose, lipid transport, and estradiol 17-beta-dehydrogenase 8 (*HSD17B8*) gene associated with estrogen biosynthetic process were highly expressed in major growth stage II group (**Figure 2A, 2B Cluster1**). Genes associated with oogenesis, border follicle cell migration, steroid metabolic process and lipid transport were highly expressed in completely sex mature precocious group enriched in cluster 2, including myosin heavy chain, non-muscle (*ZIP*), dynamin (*SHI*), Ets DNA-binding protein pokkuri (*AOP*), Protein catecholamines up (*CATSUP*), very low-density lipoprotein receptor (*VLDLR*), sulfotransferase 1A1 (*SULTLA1*), sortilin-related receptor (*SORL1*), low-density lipoprotein receptor-related protein (*LRP*) (**Figure 2A, 2B Cluster 2**). Genes enriched in cluster 3 such as Delta (24)-sterol reductase (*DHCR24*), dehydrogenase/reductase SDR family member 11 (*DHRS111*), NPC intracellular cholesterol transporter 1 (*NPC1*) were involved in cholesterol metabolic process, steroid biosynthesis were highly expressed in major growth stage I (**Figure 2A, 2B, Cluster 3**).

**Figure 2.**
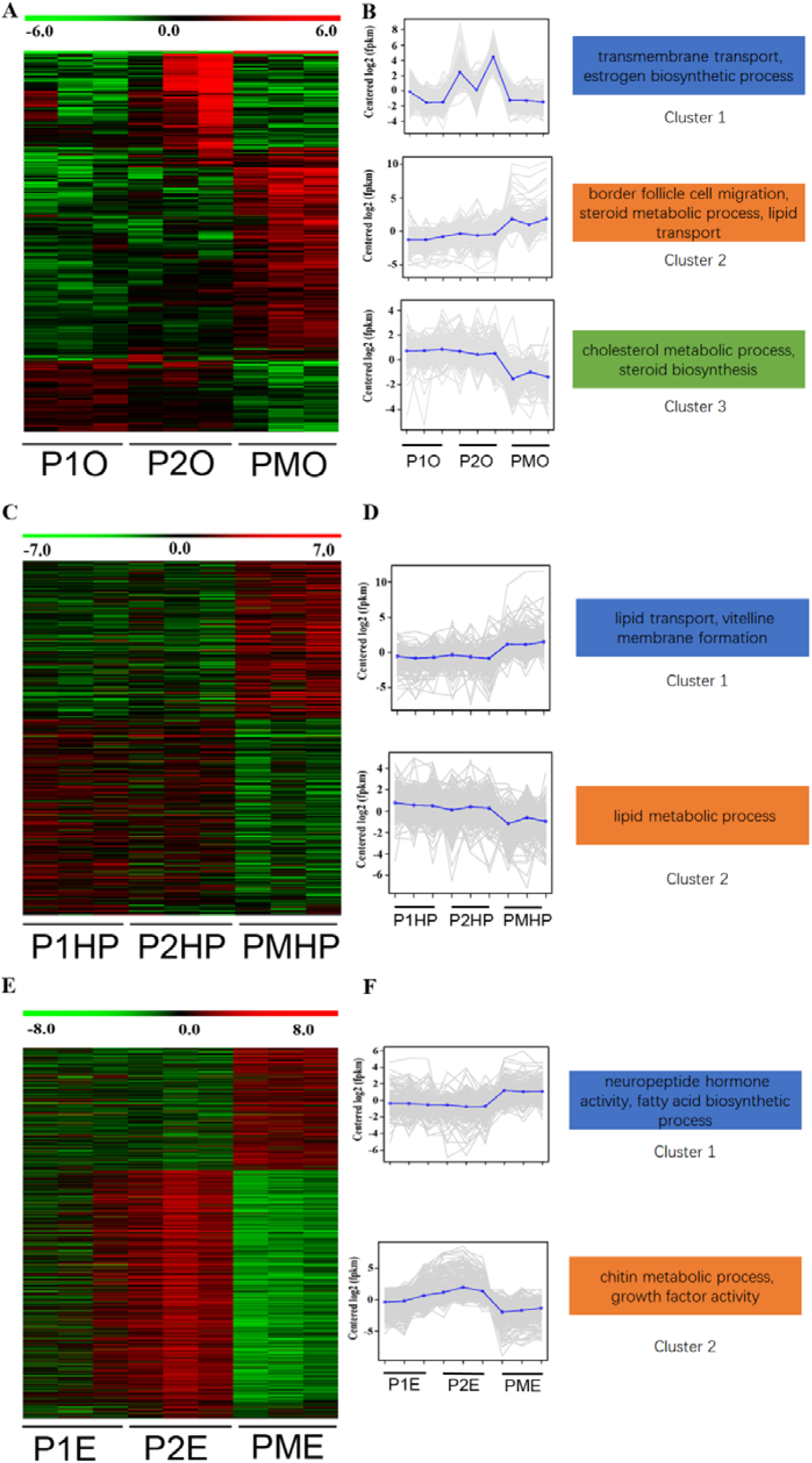
Expression patterns of differentially expressed genes in precocious ovary, hepatopancreas and eyestalk tissues. A. Heatmap of differentially expressed genes in ovary. B. Gene expression plot for the three defined clusters. C. Heatmap of differentially expressed genes in hepatopancreas. D. Gene expression plot for the two defined clusters. E. Heatmap of differentially expressed genes in eyestalk. F. Gene expression plot for the two defined clusters. P1O, P1HP, P1E indicated precocious ovary, hepatopancreas, and eyestalk with ovary in major growth stage I; P2O, P2HP, P2E indicated precocious ovary, hepatopancreas, and eyestalk with ovary in major growth stage II; PMO, PMHP, PME indicated precocious ovary, hepatopancreas, and eyestalk with ovary in sex mature stage.

Compared sex mature precocious ovary with normal two-year-old sex mature ovary, only 11 DEGs were identified, among them up-regulated genes like Neuroparsin-A (*NPAB*), Lipase 3 (*LIP3*) in precocious ovary were associated with neuropeptide hormone activity and lipid catabolic process (Table S2, Figure 3).

**Figure 3.**
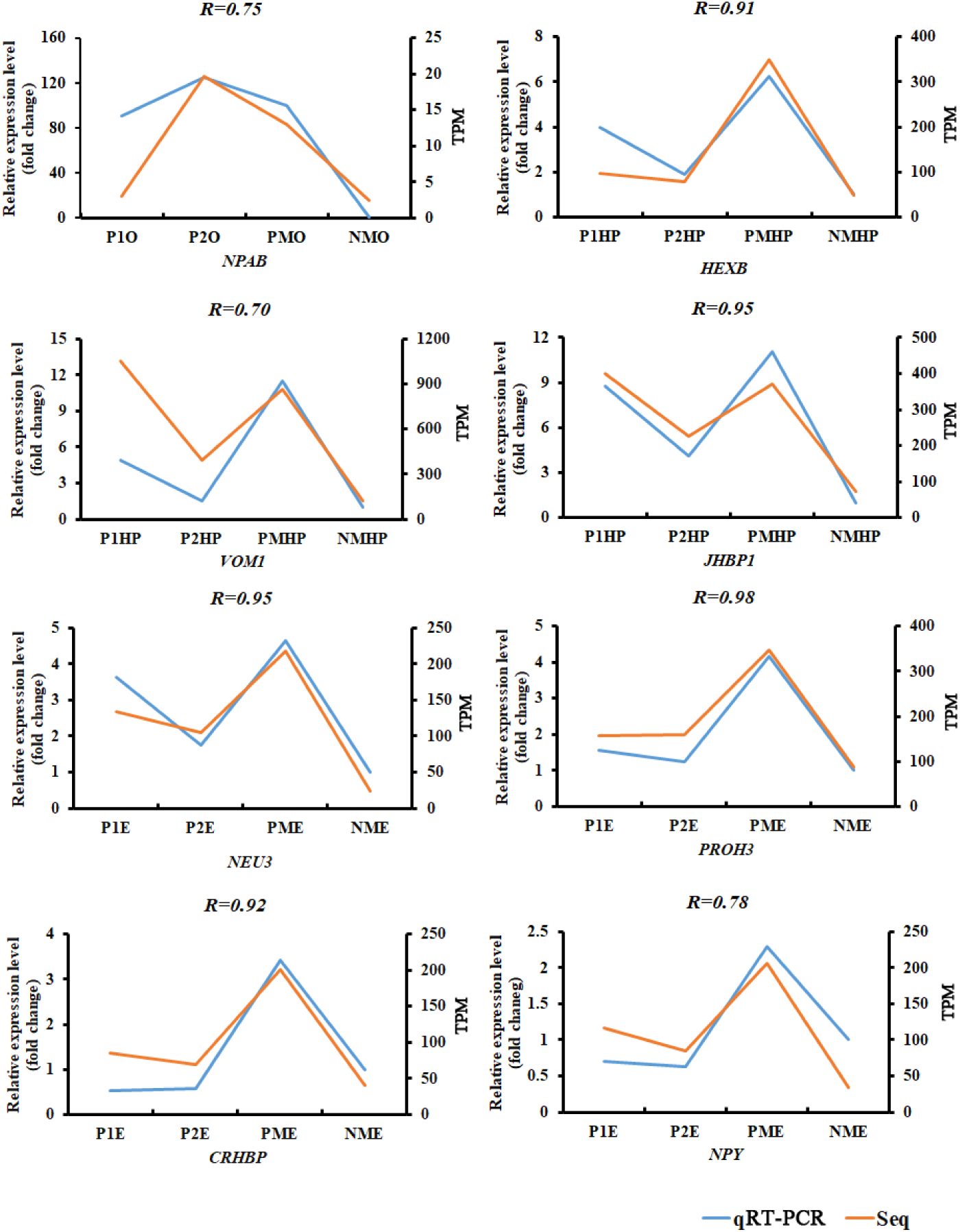
Expression profiles of eight differential expressed genes from RNA-Seq (orange) and qRT-PCR (blue) at different developmental stages of the ovary. P1O, P1HP, P1E indicated precocious ovary, hepatopancreas, and eyestalk with ovary in major growth stage I; P2O, P2HP, P2E indicated precocious ovary, hepatopancreas, and eyestalk with ovary in major growth stage II; PMO, PMHP, PME indicated precocious ovary, hepatopancreas, and eyestalk with ovary in sex mature stage.

Considering hepatopancreas tissue, a total of 806 DEGs were identified among different groups of precocious hepatopancreas. Two clusters were defined based on the hierarchical clustering results. GO enrichment indicated that genes such as ATP-binding cassette sub-family A member 3 (*ABCA3*), sodium/bile acid cotransporter (*SLC10A1*), ileal sodium/bile acid cotransporter (*SLC10A2*), vitelline membrane outer layer protein 1 (*VMO1*), vitellogenin (*VG*), beta-hexosaminidase subunit beta (*HEXB*), hemolymph juvenile hormone binding protein (*JHBP*) in cluster 1 associated with lipid transport, vitelline membrane formation, oogenesis were highly expressed in sex mature precocious group (**Figure 2C, 2D Cluster 1, Figure 3**). Genes such as glycerol-3-phosphate acyltransferase 3 (*GPAT3*), bile salt-activated lipase (*CEL*), lysosomal acid lipase (*LIPA*), pancreatic triacylglycerol lipase (*PNLIP*) in cluster 2 were associated with lipid metabolic process (**Figure 2C, 2D Cluster 2**).

Compare normal two-year-old sexual mature hepatopancreas with sexual mature precocious hepatopancreas, 372 DEGs were identified, Genes up-regulated in completely sexual mature precocious hepatopancreas like *VMO1, VG* were enriched in vitellogenin synthesis, vitelline membrane formation, lipid transport biological process (Table S2, Figure 3).

For eyestalk tissue, a total of 1,081 DEGs were identified among different groups of precocious eyestalks. Two clusters were defined based on the hierarchical clustering results revealing different expression patterns. GO enrichment analysis indicated genes such as neuropeptide F (*NPF*), *MIH, CHH*, vasotocin-neurophysin VT 1 (*VT1*), glycoprotein hormone beta-5 (*GPHB5*), Cytochrome P450 18a1 (*CYP18A1*), corticotropin-releasing factor-binding protein (*CRHBP*) in cluster 1 highly expressed in sexual mature precocious eyestalk were associated with neuropeptide hormone activity, fatty acid biosynthetic process (**Figure 2E, 2F Cluster1, Figure 3**). Genes such as cuticle protein CP498, cuticle protein AM1159, insulin-like growth factor-binding protein-related protein 1 (*IGFBP*), platelet-derived growth factor receptor alpha (*PDGFRA*), protein 60A (*GBB*) in cluster 2 were involved in chitin metabolic process, growth factor activity (**Figure 2E, 2F Cluster 2**).

Between normal two-year-old sexual mature eyestalk and sexual mature precocious eyestalk, 449 DEGs were identified, up-regulated genes like *VT1, NPF*, Prohormone-3 (*PROH3*), helicostatins (*helicostatins*), pro-neuropeptide Y (*NPY*) in completely sexual mature precocious eyestalk were associated with neuropeptide hormone activity, neuropeptide signaling pathway (Table S2, Figure 3).

### DEGs related to neuropeptide hormone activity and lipid transport

Gene expression values from a total of 15 DEGs (GO:0005184, neuropeptide hormone activity) and 12 DEGs (GO:0006869, lipid transport) from the studied individuals were extracted. 12 out of 15 DEGs related to neuropeptide hormone activity were identified in eyestalk tissue and most of the DEGs (*MIH, GPHB5, NPA, CHH, VT1, NPF, RPCH, PDH1, CCAP, NPY*) were upregulated in precocious eyestalk than in normal sex mature eyestalk. 11 out of 12 DEGs related to lipid transport were identified in hepatopancreas tissue, and most of the DEGs (*SLC10A1*, Apolipophorin, *VG, LDLR-A, SLC10A2*) were upregulated in precocious hepatopancreas than in normal sex mature hepatopancreas (Table S3).

The correlation coefficient adjacency matrix indicated that DEGs in eyestalk annotated as neuropeptide hormone activity such as *NPY, RPCH* (SP|Q23757), *GHBP5, NPF, ORCKA* (SP|Q9NL83_ORCKA), *CCAP*, Helicostatins SP|Q44314_ALLP) were positively correlated with DEGs in hepatopancrea annotated as lipid transport such as *VG* (SP|Q6RG02), *NPC2* (SP61917), *SLC10A1* (SP|Q14973), *SLC10A2* (SP|Q28727) (*P*<0.05) (Figure 4A, red shade area). Meanwhile, DEGs in hepatopancreas annotated as lipid transport such as *VG* (SP|Q6RG02), *NPC2* were positively correlated with DEGs in ovary annotated as oogenesis, border follicle cell migration, steroid metabolic process, lipid transport (Figure 4B, yellow shade area) (*P*<0.05). However, most DEGs in eyestalk annotated as neuropeptide hormone activity were not correlated with DEGs in ovary, only *SLC10A3* (SP09131) showed positively correlation with Helicostatins, *RPCH, GHBP5*, and *CCAP* genes (**Figure S1**).

**Figure 4.**
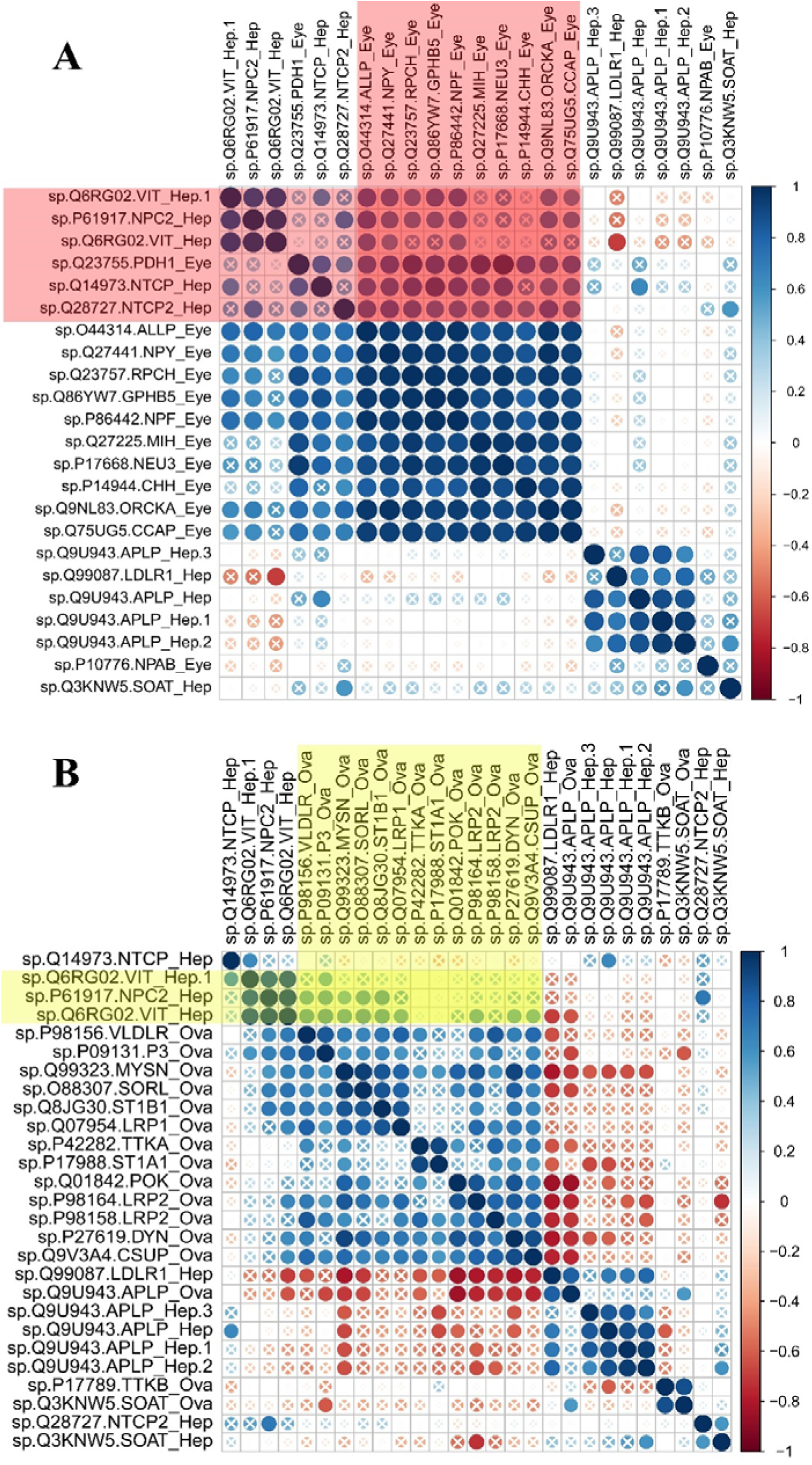
Pearson correlation of gene expression levels of DEGs between tissues (ovary, hepatopancreas, eyestalk). A. Pearson correlation of gene expression levels of DEGs between hepatopancreas and eyestalk. Red shade indicated the positively correlated genes in hepatopancreas and eyestalk (*P*<0.05). B. Pearson correlation of gene expression levels of DEGs between hepatopancreas and ovary (*P*<0.05). “X” symbol indicated P value >0.05 for the Pearson correlation.

### Discussion

Sexual precocity is a complex biological process with many genes/pathways involved in specific organs to induce gonad early development [1, 5]. In this study, we utilized the ratio of abdomen sternite length to discriminate ovary developmental stages and believed it is a convenient method for the detection of potential precocious *E. sinensis* in advance. *E. sinensis* with the ratio of abdomen sternite length above 0.70 during the first year are potential precocious *E. sinensis* that should be abandoned in the aquaculture. Compared with the expression profiles of ovary between two-year old and one-year old sex mature *E. sinensis*, only 11 DEGs were identified and the tissue histology observation showed normal function of precocious ovary which indicated that precocious *E. sinensis* are functional and able for spawning [2]. *Neuroparsin-A (NPA)* is believed to be involved in the regulation of insect reproduction through inhibit the effects of juvenile hormone was highly expressed in precocious *E. sinensis* indicating that early ovary development may due to the abnormal expression of *NPA* caused by environment factors [26]. However, comprehensive functional studies need to be conducted to reveal the molecular mechanism. Our results also provided novel insights into the genetic interaction of eyestalk and hepatopancreas on ovary early development of precocious *E. sinensis*.

Significant different gene expression profiles were identified in hepatopancreas and eyestalk between normal and precocious *E. sinensis*, which indicating the important function of hepatopancreas and eyestalk in regulating ovary development. Hepatopancreas is an essential organ for energy storage and metabolism of crustaceans, providing required energy for the growth and development [16, 27]. It is also a site for the synthesis and metabolism of certain steroid hormones required by crustaceans, which plays essential roles in vitellogenesis process [17, 28]. In this study, DEGs associated with lipid metabolic process were upregulated in major growth stage I, stage II during vitellogenesis process in precocious *E. sinensis* (Figure 2B, cluster1), indicating the initiation of ovary development depend on the lipid metabolic in hepatopancreas, which may provide energy and steroid hormones for early ovary development [29, 30].

Previous studies also pointed out that nutriment like sugar/lipid will be absorbed and accumulated in hepatopancreas, and excessive nutrition will be transferred to gonad continuously which will induce gonad early development [31]. Interestingly, DEGs associated with lipid transport were also upregulated in precocious *E. sinensis* compared with normal *E. sinensis* in both hepatopancreas and ovary tissues (Figure 2B, Cluster 2, Figure 3). These up-regulated lipid transport related genes may provide the genetic basis for the lipid transfer from hepatopancreas to ovary [30-32]. The hepatopancreas index measured in this study decreased with the increased gonad index also supported that hepatopancreas may transfer required energy/lipid to ovary (**Figure 1)**. Meanwhile, vitellogenin (VG) is the key factor component in vitellogenesis in *E. sinensis*, which include endogenous VG and exogenous VG. Endogenous VG is believed to be synthesized by oocyte, while exogenous VG is believed to be synthesized in hepatopancreas and transferred to ovary [33]. In this study, *VG* genes were significantly up-regulated in hepatopancreas of precocious *E. sinensis*, indicating hepatopancreas is the main vitellogenin synthesis site in *E. sinensis* and all the VG protein required for vitellogenesis originated from hepatopancreas. Meanwhile, *VG* genes expression in hepatopancreas was positively correlated with oogenesis, lipid transport DEGs in ovary indicating excessive VG expression in hepatopancreas may stimulate the ovary development in precocious *E. sinensis*. *VMO1* which is associated with vitelline membrane formation was also up-regulated in precocious *E. sinensis*, indicating hepatopancreas may also participate in the vitelline membrane formation process.

Our results confirmed the essential roles of hepatopancreas in regulating ovary development, energy and steroid hormone required for oogenesis; vitellogenin synthesis and vitelline membrane formation for vitellogenesis were fulfilled in hepatopancreas. The intrinsic genetic factors of sexual precocity in *E. sinensis* at some content due to the abnormal expression of the above-mentioned candidate DEGs in hepatopancreas.

It is believed that X-organ-sinus gland complex system existed in eyestalk is an important neuroendocrine system in crustaceans [7]. Previous study indicated by regulating the endocrine system of shrimp and crab, the growth rate and the mature time will be improved, and eyestalk ablation (ESA) is a way to stimulate gonad development and ovulation [34]. Consistent with previous hypothesis, more neuropeptide hormone like *NPF, NPY*, Prohormone-3 were identified as up-regulated in precocious eyestalk in this study indicating the essential regulatory mechanism of eyestalk in ovary development. *NPF* and *NPY* belong to NPY family are neuropeptide hormone that accelerates ovarian maturation in female *Schistocerca gregaria* [35], and plays a role in the regulation of visceromotor functions during egg laying [36]. *CHH, MIH* have long been considered to inhibit crustacean molting, and after *E. sinensis* finish their last reproductive molting like insects, they will stop molting and initiate their gonad development [37, 38]. Therefore, extremely highly expressed *CHH, MIH* in eyestalk may inhibit the molting and induce precocity. More interestingly, DEGs in eyestalk such as *NPY, RPCH*, Helicostatins, *GHBP5, NPF, ORCKA, CCAP* were positively correlated with *VG* genes expression in hepatopancreas which indicate these neuropeptide hormone genes may target hepatopancreas and induce *VG* expression, however, further functional experiments need to be conducted to confirm the hypothesis (Figure 4B). All the upregulated genes in precocious eyestalk indicated the neuropeptide hormone synthesized in eyestalk may stimulate ovary development like HPG axis. Our results pointed out that eyestalk is indeed essential organ for gonad development and may play essential roles in regulating vitellogenesis in precocious *E. sinensis*. However, the interplay of eyestalk, hepatopancreas, and ovary requires more further functional researches.

Sexual precocity is a serious situation in the aquaculture of *E. sinensis*, scientific researchers and farmers were struggled with the solutions and genetic mechanism of sexual precocity. In this study, there were few expression differences identified between precocious and normal sexual mature ovary, and the early development of ovary may be affected by abnormal developed eyestalk and hepatopancreas. Several stimulation factors such as high temperature, salt, stock density, nutrition may induce the metabolic disorder of genes associated with neuropeptide hormone, steroid hormone as identified in this study, leading to the abundant expression and accumulation of VG in hepatopancreas and further initiating the ovary development. However, comprehensive functional studies should be conducted to further reveal the genetic mechanism of sexual precocity, especially the regulatory mechanism for eyestalk and hepatopancreas. Our study provides valuable genetic resources for the research of sexual precocity in *E. sinensis* in future. Meanwhile, the effective convenient phenotype measurement method and related candidate DEGs identified in this study will provide guidance for the detection of precocious *E. sinensis* in aquaculture.

## Acknowledgements

This work was funded by Agriculture Research System of Shanghai, China (Grant No. 201804), Shanghai Agriculture Applied Technology Development Program, China (Grant No. G2017-02-08-00-10-F00076), Shanghai Science and Technology Committee Programs, China (No.16391905300; No. 13DZ2251800), Leading Agricultural Talents in Shanghai Project, China (Grant No. D-8004-16-0217).

